# Label-free monitoring of therapy response in 3D spheroids using lab-on-a-chip impedance spectroscopy

**DOI:** 10.1101/2025.09.08.674936

**Authors:** Gregory Macke, Maulee Sheth, Manju Sharma, Supasek Kongsomros, Maria Lehn, Trisha M. Wise-Draper, Vinita Takiar, Leyla Esfandiari

**Affiliations:** Department of Biomedical Engineering, University of Cincinnati, Cincinnati, Ohio 45221, USA; Division of Hematology/Oncology, Department of Internal Medicine, University of Cincinnati College of Medicine, Cincinnati, Ohio 45219, USA; University of Cincinnati Cancer Center, Cincinnati, Ohio 45267, USA; Department of Radiation Oncology, University of Cincinnati College of Medicine, Cincinnati, Ohio 45219, USA; Department of Electrical Engineering and Computer Science, University of Cincinnati, Cincinnati, Ohio 45221, USA; Department of Environmental and Public Health Sciences, University of Cincinnati, Cincinnati, Ohio 45267, USA

## Abstract

The high incidence and mortality of cancer continue to drive research and development of effective therapies. Lab-on-a-Chip (LOC) platforms have emerged as powerful alternatives to traditional biological evaluation methods, offering reduced complexity, lower costs, and improved throughput. In parallel, the integration of non-invasive and non-destructive sensing techniques have expanded opportunities for real-time and label-free analysis. Electrical impedance spectroscopy (EIS), which exploits the intrinsic dielectric properties of cells, has shown promise for the quantitative evaluation of 3D cellular structures. In this study, we demonstrate the application of LOC-based EIS to assess the bioeffects of radiotherapy on 3D head and neck cancer spheroids. Our results establish EIS as a viable tool for real-time monitoring of treatment-induced changes in 3D tumor models, supporting its potential in preclinical cancer research and therapeutic screening.

## Introduction

Cancer is the second leading cause of death in the United States, with millions of new cases projected in 2025, and its high incidence and mortality have driven the development and screening of targeted therapies that improved the overall 5-year survival rate. ^1,2^ These advanced interventions employ strategies such as pharmaceutical treatments, radiotherapy, and gene therapies that are initially developed through in vitro screening, establishing efficacy and safety across cellular and subsequently, animal investigations. ^3^ However, the barriers to entry of these essential studies stifle additional progress toward clinical applications. Gold standard in vitro biological evaluations commonly involve techniques that rely on staining cells or lysing them for characterization. These methods often require skilled personnel and involve time-consuming procedures, need expensive equipment, and are destructive to the sample being tested. ^4^

To overcome these limitations, there is an increasing interest in lab-on-a-chip (LOC) platforms that consume smaller reagent and sample volumes, reduce costs, and improve accessibility with streamlined protocols. ^5-7^ The innovation of LOCs integrated with noninvasive, high-throughput, and label-free sensing technologies is capable of mitigating challenges associated with the screening of novel therapies. A variety of noninvasive sensing techniques have been developed to detect different physical or biomolecular properties of pathological cells and tissues. ^8, 9^ Leveraging changes in the electrical characteristics of biological samples through electrical impedance spectroscopy (EIS) has an established basis of efficacy for non-invasive, label-free sensing. An on-chip application of this technology could provide a promising candidate for quantified in vitro assessment of physiologically relevant 3D cell structures such as spheroids and organoids. ^10^

Since its conception, EIS capabilities have been extended toward biomedical applications, where it has been applied to in vitro studies aiming to monitor the activity of pharmaceuticals and other treatments. ^11^ During EIS measurement, an AC electric field can be applied at an individual frequency to detect a specific phenomenon or swept over a broad frequency range to measure different biologically relevant aspects of the sample across the spectrum. Studies have adopted lower frequencies in the alpha dispersion, ranging from approximately 10 Hz to 10 kHz in cells, to monitor proliferation and viability impacts on monolayer and hydrogel cultures in response to anti-cancer pharmaceuticals. ^12, 13^ Within this dispersion, the impedance signal is expected to be sensitive to surface charges and membrane relaxation, but it can also be strongly influenced by electrode polarization. ^12, 14^ At higher frequency ranges in the beta dispersion, ranging from approximately 10 kHz to 10 MHz, the electric field can penetrate the cell and has been utilized to discern spheroids from microspheres as well as to evaluate viability impacts using opacity ratio. ^12, 14-16^ The impedance signal within this dispersion is expected to primarily be influenced by the cellular membrane and cytosolic contents. ^12, 14^

Physical and biomolecular cargo alterations within cells in response to an applied cancer therapy can lead to measurable differences in their dielectric properties across these dispersions, creating the potential to noninvasively quantify therapeutic effect in 3D cell structures. EIS has been previously employed to detect the cytotoxic effects of pharmaceuticals and the cellular activity of stem cell differentiation in spheroids and has the potential to facilitate the development and testing of further targeted therapies. ^16, 17^ In this study, we engineered a reversibly sealed 3D printed LOC with integrated sensing electrodes to measure the impedance of head and neck cancer (HNC) spheroids. The overall size and radiation-induced cellular changes were found to influence the dielectric properties of Cal27 oral squamous cell carcinoma spheroids. These results demonstrate that the LOC implementation of EIS can be used to noninvasively discern the effects of radiotherapy within 3D HNC spheroids in a label-free manner. The LOC-EIS integration has the potential to be further expanded for investigation aimed toward high-throughput drug screening and personalized medicine approaches.

### Experimental Materials and Methods

#### Cell growth and maintenance

Oral squamous cell carcinoma Cal27 cells (ATCC, Manassas, VA, USA) were grown in accordance with previously published work from our group, as briefly described. ^18^ Cells were cultured on surface-treated tissue culture plates (Fisher Scientific, Waltham, MA) in DMEM (Corning, San Diego, CA) containing 10% exosome-depleted fetal bovine serum (Fisher Scientific, Waltham, MA), 1% penicillin-streptomycin (Fisher Scientific, Waltham, MA), 1% non-essential amino acids (Fisher Scientific, Waltham, MA), 1% sodium pyruvate (Fisher Scientific), and 1% L-glutamine (Fisher Scientific, Waltham, MA). The cells were incubated using a Heracell Vios 160i CO2 Incubator (Fisher Scientific, Waltham, MA) at 37 °C with 5% CO2. Cultures were allowed to proliferate until they reached approximately 70% confluency, and after at least 2 passages in exosome-depleted culture media, cells were seeded into spheroids.

#### Spheroid culture

Spheroids were cultured in low-adherence Nunclon Sphera-Treated U-bottom microplates (Fisher Scientific, Waltham, MA) following an adapted seeding protocol. ^19^ Briefly, cells cultured in the tissue culture plate were trypsinized and prepared into a cell suspension. 100 μl of the cell suspension was seeded into individual U-bottom wells at the desired cell density. The microplates were then centrifuged using an Allegra X-30R (Beckman Coulter, Brea, CA) at 500 relative centrifugal force (rcf) for 5 minutes. The wells were visually inspected to ensure that the cells had collected at the bottom. During incubation, the collected cells proliferated and grew together into 3D spheroid structures. The culture media was refreshed every 48 hours by removing and replenishing 65 μl of the total 100 μl volume. Spheroids were imaged on days 2, 4, and 7 prior to media changes using an Eclipse TE2000-5 microscope (Nikon, Tokyo, Japan) equipped with a Zyla CMOS camera (Andor, Belfast, Ireland) and NIS-Elements software (Nikon, Tokyo, Japan).

#### Microfluidic lab-on-a-chip assembly

The lab-on-a-chip was fabricated using a Form 3 Stereolithography (SLA) 3D-printer (Formlabs, Somerville, MA) in clear resin. The SLA-printed channel [Figure 1a] was spray-coated with a dry Teflon lubricant (WD-40 Company, San Diego, California) to prevent spheroids from adhering to the channel walls. To assemble the device, the electrodes were placed into slots along the wings of the chip. The lengths of the stainless-steel bodies sit in the grooves, leaving the pins exposed for connection to external equipment. Channels holding the electrodes were designed to have precise tolerances to ensure accurate positioning and minimize fluid leaks. A PDMS sheet was placed over the top of the channel and the electrodes to serve as a compliant closure. The assembly was aligned on the bottom glass slide and placed between two 3D-printed compression plates. The SLA channel was then compressed against a PDMS sheet using small bolts and wing nuts [Figure 1b], creating a reversible seal for easy assembly and reusability. ^20^ This set up also allowed for manual control of the applied pressure and microscope measurement. ^21^ The assembly was flipped 180° to visualize the spheroid as it was washed, aligned, and impedance measurements were taken.

**Figure 1.**
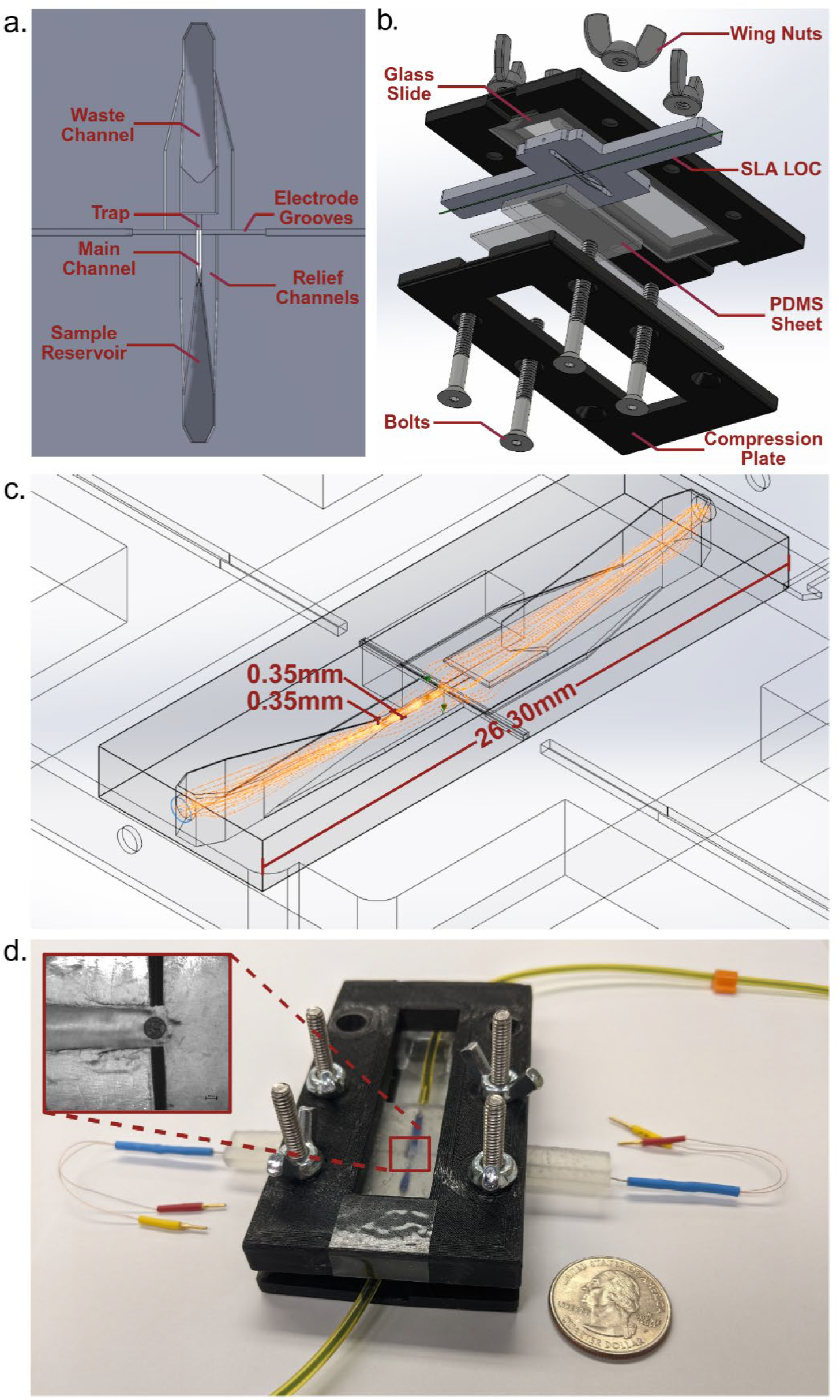
Microfluidic lab-on-a-chip for spheroid entrapment and impedance measurements. (a) 3D CAD model of the microfluidic channel geometry. (b) Exploded view depicting different parts of the CAD model assembly. (c) Simulated fluid flow through the microfluidic channel. (d) The assembled prototype with colored fluid to show the channel’s path. Callout shows a 200 μm carboxyl polystyrene microsphere trapped and aligned between the sensing electrodes (10X), scale bar = 100 μm.

#### Spheroid trapping

Spheroids were aspirated in 2 μl of media and inserted into the sample reservoir near the front of the channel. Following assembly of the device, 1X PBS filtered with a 0.02 μm inorganic membrane filter (Cytiva, Marlborough, MA) was pumped into the fluid inlet at 50 μL/min using a syringe pump (New Era Pump Systems, Inc.) [Figure 1c]. The buffer solution was flown through the channel for five minutes, during which it mixed with the culture media and carried the spheroid with the current. The channel tapered to an electrode gap size of 350 μm, guiding the spheroid to be trapped due to a decrease in channel height between the electrodes. The surface of the trapping construct that is in contact with the sample was designed to be a flat face to reduce damage caused by sharp corners. Shallow and wide relief channels allow fluid to continue flowing around the spheroid to prevent clogging and reduce pressure. The fluid flow washed away and replaced by the culture media with 1X PBS allowing for measurement of the aligned spheroid [Figure 1d]. At the trapping zone, the channel widened to reduce the fluid pressure and direct flow out of the channel. The assembly was disassembled, washed with isopropyl alcohol (Sigma-Aldrich, St. Louis, MO), dried with compressed air (Digital Innovations, Bellingham, WA), and the inlet hose was bled of air bubbles before the next use.

#### LabOne data acquisition and post processing steps

Impedance measurements were performed with a 4-terminal arrangement utilizing a cluster microelectrode (FHC, Bowdoin, ME) with 50 μm poles on either side of the spheroid, driven and recorded by an HF2LI lock-in amplifier (Zurich Instruments, Zurich, Switzerland) and an HF2TA current amplifier (Zurich Instruments, Zurich, Switzerland). Experimental settings and data acquisition were controlled through the associated LabOne software (Zurich Instruments, Zurich, Switzerland). An AC 100mV peak probing voltage was applied. Each data point recorded on the waveform represented an average of 10 measurements. The waveform consisted of 1000 points distributed over a logarithmic scale from 1 kHz to 10 MHz. A voltage representation of the magnitude and phase for the current and differential voltage measurements were recorded. All input channels were set to DC coupling. The transimpedance gain was set to 1 kV/A, and the voltage gain was set to 1 to create a suitable signal to noise ratio. The current measurement was taken with a 50 Ω input impedance, and the differential voltage measurement was taken with a 1 MΩ input impedance to reduce current leakage. The unwrap phase function was enabled for both measurement channels to reduce discontinuities, and the recorded data was exported from the LabOne software. Waveform data were loaded into a custom MATLAB script for digital signal processing and statistical analysis. This signal processing was adapted from programs used in previous studies conducted by our research group. ^22, 23^ The magnitude and phase waveforms were converted into complex numbers. Current waveform amplitudes were scaled by a factor of 2 to account for the reduction caused by the 50 Ω input impedance. The measured current waveform was converted to amps according to the current amplifier manufacturer’s specified transfer function. The impedance of the spheroid was then calculated using Ohm’s law from the current and differential voltage measurements. To compare measured impedance across experimental groups at discrete frequencies, a 15-point running average filter was applied to the waveforms. A polynomial approximation was performed using the polyfit function to estimate value at specific frequencies of interest for statistical analysis.

#### Sized-based evaluation

Spheroids were seeded at 2000 and 4000 cells per well and grown for 7 days to maximize diameter disparity. On day 7, individual spheroids were aspirated and their impedance measured to detect dielectric property changes resulting from spheroid size differences. The optical size of the aligned spheroids was measured immediately following impedance measurements, and the areas were calculated for comparison using an elliptical estimation. The elliptical area estimations were calculated by measuring the vertical and horizontal diameters of each spheroid, and the corresponding radii were used to calculate the area of the ellipse.

#### Spheroid irradiation and cellular responses

On the first day following seeding, 160 kV X-ray radiation was applied at dose of 4 Gy using a Cell Rad+ (Precision X-Ray Irradiation, Madison, CT). ^24^ On day 4 of growth, individual samples from control (untreated) and irradiated spheroids were aspirated, and their impedance was measured using the LOC device. ^24^ To assess the radiation effect in cellular level, the protein levels of γ-H2AX, Ki-67, and cleaved caspase-3 were measured by western blot using a protocol adapted from previous experiments conducted within our research group. ^25, 26^ Spheroids were harvested on day 1 of growth, 3 hours post-radiation exposure, to compare the presence of γ-H2AX among experimental groups. Additional spheroids were grown to day 4 and harvested to detect the levels of Ki67 and cleaved caspase-3, as well as cellular metabolic activity, at the time of impedance testing.

#### Western blot

For western blotting, 45 spheroids were combined and centrifuged at 500 ×g for 5 minutes, forming a pellet at the bottom of the microtube. The supernatant was removed and stored at -80 °C. The spheroid pellet was washed 3 times in ice-cold 1XPBS and resuspended in 50 μl of RIPA buffer plus protein inhibitor. The lysate was mixed and then sonicated in an ultrasonic bath (VWR Symphony, Radnor, PA) with cold water for 3 cycles of 1 minute active and 1 minute of rest. ^27^ The suspension was then incubated on ice for an additional 10 minutes.

Protein (20 μg) was acquired from the lysate and combined with 4X Leammli buffer (Bio-Rad, Hercules, CA) to prepare the mixture for gel electrophoresis. After a 5-minute incubation at 95°C, the preparation was run in the NuPAGE 12%, Bis-Tris, 1.0mm, Mini-PROTEAN TGX precast gels (Bio-Rad, Hercules, CA) for 50 minutes and blotted onto a PVDF membrane with a Trans-Blot Turbo System (Bio-Rad, Hercules, CA). The membrane was blocked with EveryBlot blocking buffer (Bio-Rad, Hercules, CA) and then incubated with 1:1000 anti-gamma H2A.X (Abcam, Waltham, MA), anti-Ki67 (Abcam, Waltham, MA), and anti-cleaved caspase-3 antibodies (Abcam, Waltham, MA) overnight at 4°C. The membranes were washed and labeled with 1:1000 goat anti-rabbit (Abcam, Waltham, MA) and goat anti-mouse (Abcam, Waltham, MA) secondary antibodies as appropriate, by incubating for 1 hour at 25°C. The detected signal was developed with Clarity Western ECL Substrate (Bio-Rad, Hercules, CA) and captured using the ChemiDoc MP Imaging system (Bio-Rad, Hercules, CA). Band intensities were quantified using ImageJ softwere (NIH, USA) and normalized to β-actin levels for statistical analysis. ^28^

#### Cell Viability assay

MTT (3-(4,5-dimethylthiazol-2-yl)-2,5-diphenyltetrazolium bromide) colorimetric assays were performed using a CyQUANT MTT Cell Viability Assay Kit (Invitrogen, Waltham, MA) adapted from the manufacturer’s protocols, briefly described as follows. For each experimental group, 6 spheroids were centrifuged at 500 ×g for 5 minutes and resuspended in 100 μl of TrypLE (Gibco, Waltham, MA) to dissociate the spheroids into a single-cell suspension. The resulting suspension was centrifuged again at 500 ×g for 5 minutes and resuspended in a combination of 100 μl of fresh culture media and 10 μl of 12-mM MTT stock solution. A blank mixture of culture media and MTT stock solution without cells was prepared for baseline comparison. The cells were incubated in the solution for 2 hours at 37 °C. Following incubation, the cell suspension was centrifuged and 85 μl of the supernatant was removed. 50 μl of DMSO (Fisher BioReagents, Waltham, MA) was added and mixed thoroughly. After a 10-minute incubation, the entire 75 μl volume was transferred into a flat-bottom microplate (Thermo Scientific, Waltham, MA), and the absorbance was measured at 540nm using a BioTek 800 TS microplate reader (Agilent, Santa Clara, CA).

#### Statistical analysis

The impedance of 5 replicate spheroids was measured for each experimental group, and the 3 spheroid spectra with the smallest standard deviation over the frequency range of interest were taken for statistical analysis. Statistical testing was performed using GraphPad Prism 9 (GraphPad, San Diego, CA). Each experiment was independently repeated 3 times, and the results were tabulated to observe reproducibly significant differences.

Data points were represented as the mean ± standard deviation. An unpaired t-test was performed on the impedance or opacity magnitude of 3 samples to compare their mean at individual frequencies of interest. A Gaussian distribution was assumed based on normality testing. When the corresponding F-test yielded a p-value greater than 0.05, equal standard deviations between samples were assumed. If the F-test proved statistically significant, a t-test with Welch’s correction was performed to account for unequal standard deviations. Each evaluation was conducted as a 2-tailed test with a 95% confidence interval and p-value threshold of 0.05.

## Results

### Spheroids size evaluation by the EIS

To evaluate the performance of impedance measurement system in detecting differences in spheroid size, spheroids with a starting density of 2000 and 4000 cells were cultured for 7 days [Figure 2a-i]. ^29^ On day 7, spheroids were harvested, and each spheroid was aligned between the sensing electrodes of the LOC for impedance measurement [Figure 2a-ii]. A significant increase in area was observed in spheroids seeded at 4000 cells compared to those seeded at 2000 cells [Figure 2b]. Spheroids seeded at 2000 cells had an average area of 0.0446 ± 0.0058 mm^2^ with an average maximum diameter of 250.29 ± 8.05 μm. Spheroids seeded at 4000 cells grew larger, with an average area of 0.0708 ± 0.0065 mm^2^ and an average maximum diameter of 310.80 ± 20.20 μm. Analysis of the impedance magnitude from 10 kHz to 10 MHz spectra indicated a more prominent increase in impedance at lower range frequency of 10 kHz to 100 kHz for detection of the spheroid size[Figure 2c]. Further examination at discrete frequencies across 6 regular intervals ranging from 10kHz to 100kHz, revealed a significant increase in impedance between 20 kHz and 100 kHz [Figure 2d]. The average difference in impedance magnitude was 1.043 ±.286 kΩ, with a maximum increase of 1.34-fold at 20 kHz. Experiments were repeated in triplicate, and recurring significant differences were observed at 3 discrete frequencies from 60 kHz to 100 kHz [Supplementary Table 1].

**Figure 2.**
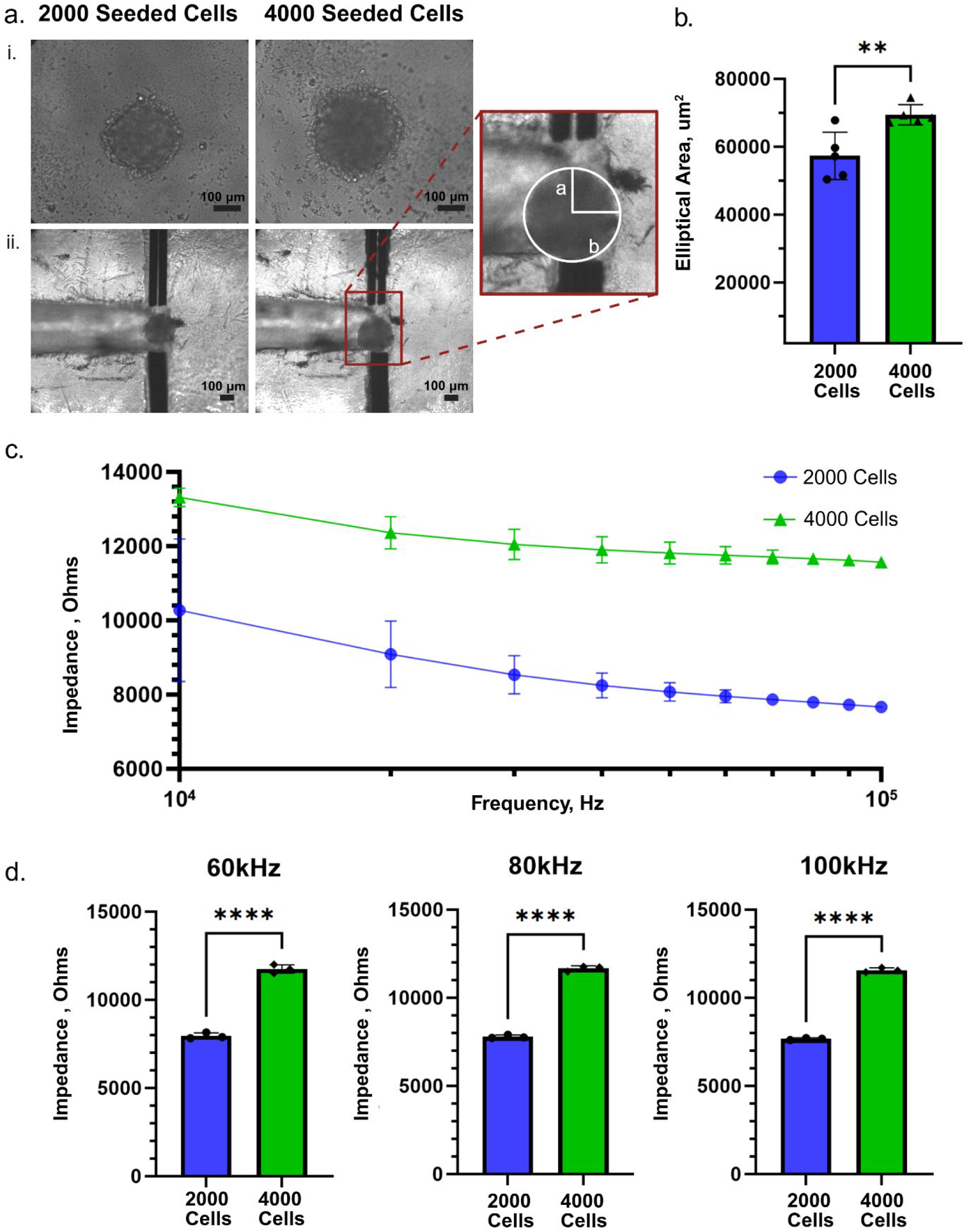
Distinction of spheroid size at low-frequency impedance measurement. (a-i) Bright field growth images of spheroids seeded with 2000 and 4000 cells (20X), scale bar = 100 μm. (a-ii) Brightfield images of aligned spheroids in the microfluidic channel (10X), scale bar = 100 μm. Callout depicts a spheroid with elliptical radii overlaid for area estimation. (b) Comparison of spheroid area estimations with varying seeding densities, n = 5. (c) Measured impedance spectra showing the mean and SD from 10 kHz to 100 kHz with varying seeding densities, n = 3. (d) Comparison of measured impedance at discrete low frequencies, n = 3. Statistical comparisons were performed using an unpaired two-tailed t-test: ^**^p < 0.005, ^****^p < 0.0001; ns, not significant.

### Mitigating size and alignment influence with opacity normalization

To reduce the influence of the sample’s size and position between the electrodes on the measured impedance, the signal can be normalized as the opacity ratio. ^30^ The opacity ratio is calculated by dividing the amplitude of impedance at each frequency Z(*f*), by the amplitude of impedance at the size-dependent reference frequency Z(*f*_ref_) (60 kHz) ^16^, shown in Equation 1. ^23^

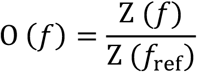

The measured impedance of spheroids seeded at a starting density of 2000 and 4000 cells were normalized as opacity ratio [Figure 3a]. The resulting opacity magnitudes were compared to determine if the electrical properties of the otherwise alike cells within spheroids with differing diameters remained consistent. When the impedance spectra of the experimental groups were compared, significant differences were observed most predominantly between 10 kHz and 100 kHz, with smaller impedance differences continuing beyond 300 kHz [Figure 3b]. Upon normalization [Figure 3c], the corresponding opacity magnitudes were not significantly different at discrete frequencies between 400 kHz to 2 MHz [Figure 3d]. This normalization revealed equivalent dielectric properties when the size dependency on impedance was compensated for in the spheroids of differing diameters.

**Figure 3.**
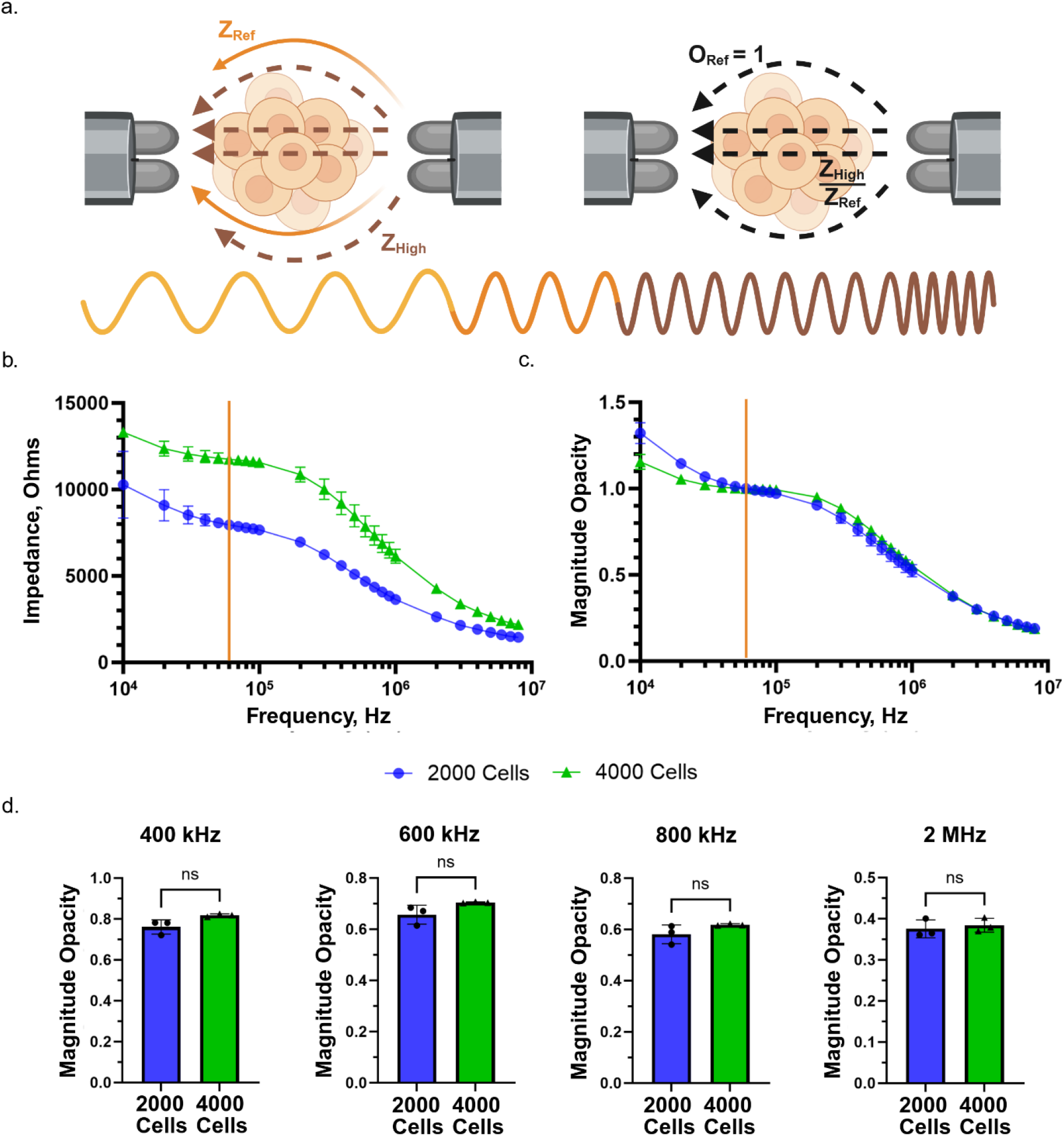
Opacity ratio in spheroids. (a) A graphical representation of the magnitude opacity normalization. ORef refers to the opacity magnitude at the size-dependent reference frequency, ZRef refers to the impedance at the size-dependent reference frequency, and ZHigh refers to the impedance at high frequency. (b) Impedance spectra of spheroids seeded using 2000 and 4000 cells, n = 3. (c) The opacity spectra of the 2000 and 4000 seeded cell spheroid groups, n = 3 spheroids. (d) Magnitude opacity evaluated at discrete frequency, n = 3. Statistical comparisons were performed using an unpaired two-tailed t-test: ns, not significant.

### Evaluation of irradiation effect on Spheroids by the EIS

To evaluate changes in dielectric properties induced by the irradiation of 3D spheroids, the opacity magnitude of control and irradiated spheroids [Figure 4a-i] was compared 3 days after exposure. Spheroids were washed and aligned between the sensing electrodes of the LOC device [Figure 4a-ii]. Their impedance was measured and converted to opacity magnitude using a size-dependent reference frequency of 60 kHz. ^16^ Comparison of the opacity magnitude spectra across a broad frequency spectrum indicated increased opacity in irradiated spheroids from 200 kHz to 3 MHz [Figure 4b]. Further examination at discrete frequencies revealed significant differences in magnitude opacity from 200 kHz to 3 MHz [Figure 4c]. The average difference in opacity magnitude was 0.109 ± 0.0270, with a maximum 1.36-fold increase in irradiated spheroids at 1 MHz. Investigations were repeated in 3 independent experiments, with recurring significant differences found at eight frequencies evaluated from 400 kHz to 900 kHz, and from 2 MHz to 3 MHz [Supplementary Table 2].

**Figure 4.**
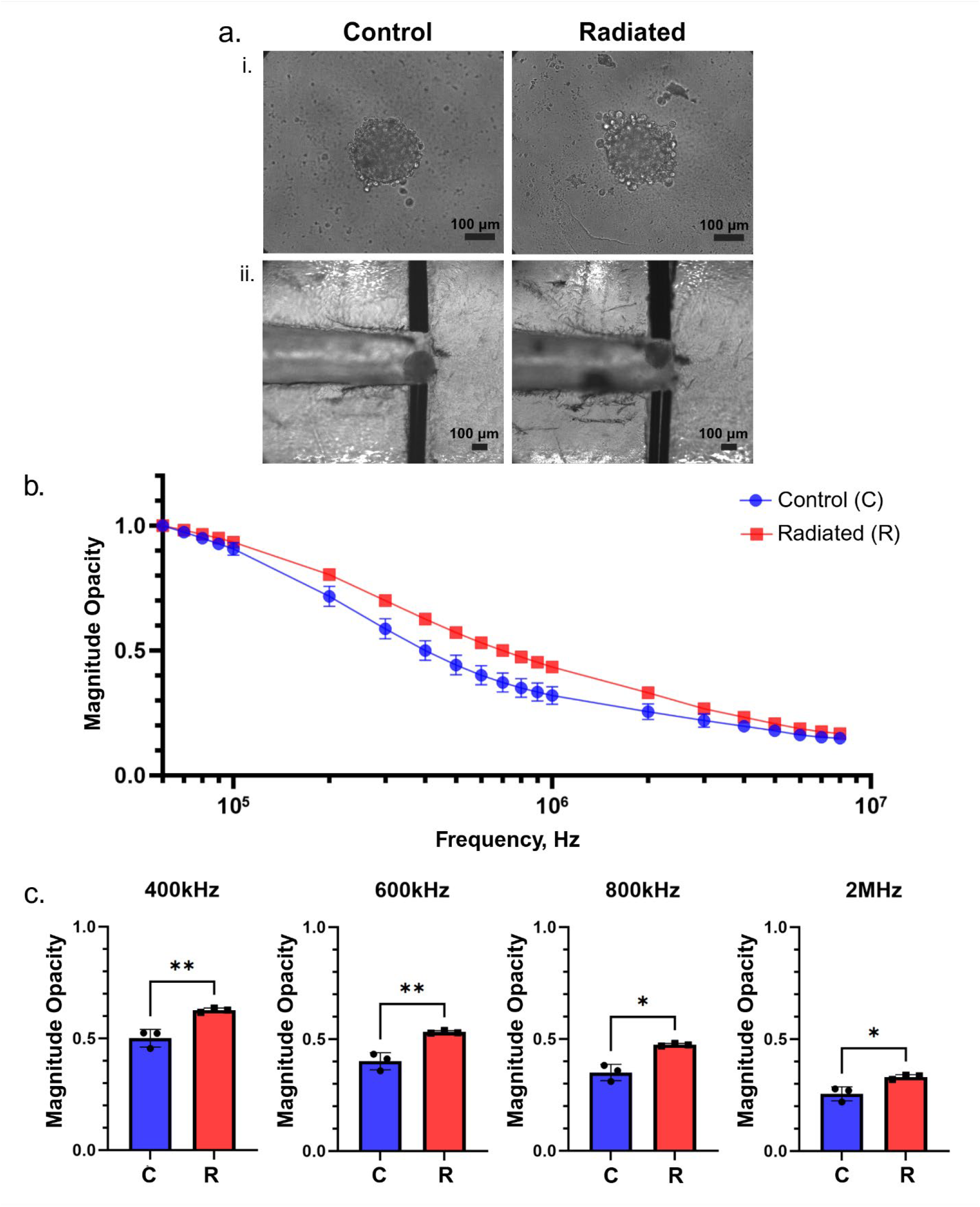
Impedance measurements of 3D spheroids 3 days after radiation treatment. (a-i) Brightfield images of control and irradiated spheroids growing in u-bottom microplate wells on day 4 of growth (20X). (a-ii) brightfield images of control and irradiated spheroids aligned between the sensing electrodes within the microfluidic channel (10X), scale bars = 100 μm. (b) The magnitude opacity spectra of control and irradiated spheroids from 60 kHz to 8 MHz, n = 3 spheroids. (c) Magnitude opacity evaluated at discrete frequencies for statistical analysis, n = 3. Statistical comparisons were performed using an unpaired two-tailed t-test: ^*^p < 0.05, ^**^p < 0.005.

### Biomolecular assessment of spheroid response to radiation

To investigate the effects of radiotherapy that can influence the changes observed in the measured opacity, spheroids were tested for relevant protein markers. Western blot analysis indicated increased protein expression of γ-H2AX, a hallmark of DNA damage, in the irradiated groups 3 hours post-exposure [Figure 5a]. Cleaved caspase-3 expression, an indicator of apoptosis, also increased, while a decrease was observed in Ki-67, reflecting reduced proliferation within irradiated spheroids at the time of impedance measurements [Figure 5b]. Actin expression was measured as a loading control. Quantification of western blot intensities demonstrated an increase to 1.35 ± 0.12 in average relative expression of γ-H2AX levels, a reduction to 0.54 ± 0.084 for Ki67, and an increase to 1.31 ± 0.11 for cleaved caspase-3 in irradiated spheroids compared to controls [Figure 5d]. Viability differences were corroborated using an MTT assay, where a significant decrease in metabolic activity was observed with an average relative expression of 84.51 ± 7.90% in irradiated spheroids compared to controls [Figure 5c].

**Figure 5.**
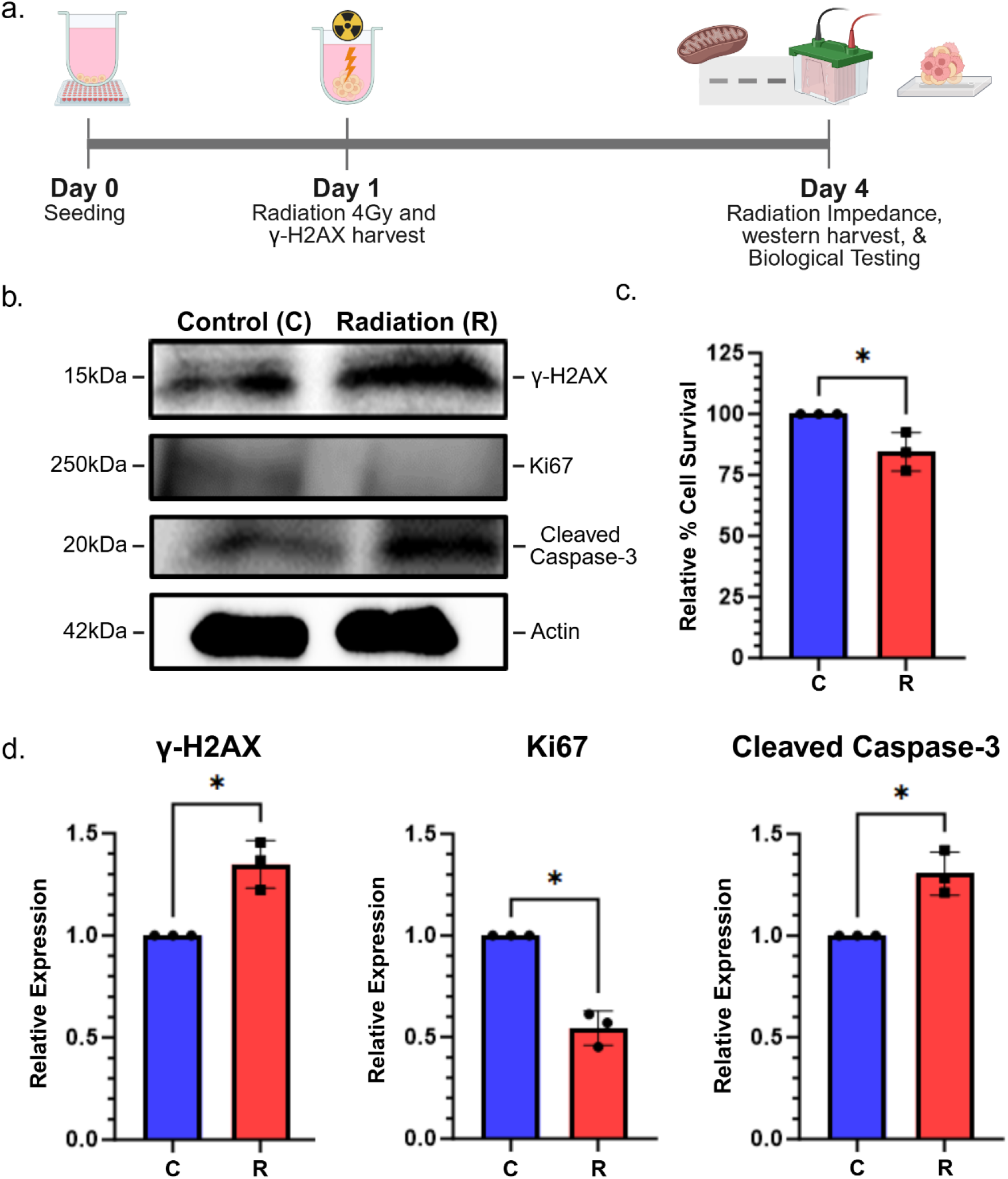
Cellular influences of radiotherapy leading to changing dielectric properties in treated spheroids. (a) Timeline of spheroid harvest for biomarkers detection. (b) Representative western blot of selected biomarkers (γ-H2AX, Ki-67, cleaved caspase-3, and actin) impacted by radiation, altering the spheroids’ dielectric properties. (c) relative cell survival between control and irradiated spheroids measured using MTT assay, n=3. (d) Quantified relative protein expression of immunoblots, n=3. (the full-length blot images were available in Supplementary figure 1).Statistical comparisons were performed using an unpaired two-tailed t-test: ^*^p < 0.05, ^**^p < 0.005.

## Discussion

In this study, we developed and utilized an integrated LOC-EIS platform to measure changes in the electrical impedance of 3D HNC spheroids in response to radiation therapy. Spheroids with smaller areas were found to have a decreased impedance magnitude at lower frequencies from 60 kHz to 100 kHz. This corroborates previous findings where spheroids with smaller optical diameters were found to have lower impedance magnitudes at 60 kHz, exhibiting moderate variability with shape, uniformity, and orientation. ^16^ While this trend has been previously reported over a limited number of discrete frequencies, our study expands the analysis across a broader frequency range with finer intervals to provide critical insight into cellular behavior in response to radiotherapy.

To account for size-and positioning-dependent variations in impedance measurements, signals were normalized using the opacity ratio. The difference in impedance magnitude between the spheroids seeded with 2000 and 4000 cells was most prevalent across lower frequencies, and to a lesser extent, present at higher frequencies. These findings are consistent with reported studies in which simulated results of 2D cell clusters with increasing cell layers, which mimic increases in diameter. ^16^ In our study, impedance measurements were normalized using the opacity ratio at a size-dependent reference frequency of 60 kHz to mitigate the influence of spheroid size and positioning. The 60 kHz reference frequency was chosen due to its demonstrated relationship with size and its prior use in evaluating cellular viability within spheroids. ^16^ The insignificant differences observed in the opacity ratio of spheroids with differing sizes indicate similar dielectric properties in the cells within, independent of the spheroid’s overall diameter. The normalization as such enables subtle changes in dielectric properties to be discerned independent of variations in spheroid size and positioning.

We next utilized our LOC-EIS platform to evaluate therapeutic response using impedance spectroscopy at higher frequency range. As radiation takes effect, cells within a spheroid begin to die, leading to a strong correlation between reduced cell viability and decreased overall spheroid diameter. ^31^ The diminishing size would be detectable through its resulting size-dependent impedance magnitude in the low-frequency range of 60 to 100 kHz. Conversely, if the spheroid continues to grow due to an insufficient dose of radiation or a resurgence, it would correspond to a different impedance magnitude in the same low-frequency range. This relationship between cellular viability and spheroid size provides a valuable application for low-frequency impedance monitoring in detecting subtle changes in spheroid phenotype. The dielectric properties of irradiated spheroids were also evaluated using the opacity ratio to minimize size-dependent errors. Irradiated spheroids showed significant increases in opacity from 400 to 900 kHz and from 2 to 3 MHz, corresponding to the beta dispersion range. Such an increase in opacity is consistent with previous studies reporting increased opacity in heat-treated spheroids and altered spheroid impedance in response to drug treatment. ^16, 32^

An increased frequency in the beta dispersion range enables the field to permeate the cells and engage with the cellular membrane and charged contents that are altered during repair and apoptosis in the spheroid. ^12, 14^ To correlate the radiation-induced increase in opacity to such changes, we performed western blot analysis on key biomolecular indicators of radiation effects. Cell concentration has been previously identified as a key contributing factor altering 3D culture impedance. ^13^ In our study, irradiated spheroids were found to have a lower expression of Ki-67 as compared to controls, indicative of reduced proliferation in response to therapy. Reduced cell proliferation within irradiated spheroids leads to fewer cells within the cluster, altering the total resistive and capacitive properties, influencing the impedance. These alterations in electrical properties, in turn, likely contribute to the detected increase in the opacity between our experimental groups.

At a cellular level, radiation-induced DNA damage triggers the phosphorylation of γ-H2AX shortly after radiation, indicative of the presence and extent of double-stranded breaks. ^33, 34^ We observed increased expression of γ-H2AX in irradiated spheroids, verifying radiation-induced DNA damage within the cells and providing an example of radiation-induced changes in charged cellular contents contributing to the detected increase in magnitude opacity. ^23, 35^ This corroborates previous impedance studies that have attributed measured changes in opacity to differences in charged molecules located within the membrane and lumen. ^23^ Additionally, we also observed an increased expression of cleaved caspase-3 within irradiated spheroids, indicative of increased apoptosis. Viability differences were also corroborated using an MTT assay, which showed a significant reduction in metabolic activity in irradiated spheroids. The compromised membranes of dying cells within spheroids, which are not cleared in in vitro platforms, have been reported to lead to higher membrane conductivity and increased opacity ratios due to swelling and membrane lysis. ^36 16^ The increased opacity of our irradiated groups can, as such, also be attributed to increased apoptosis wherein membrane rupture reduces cellular impedance by increasing conductivity, disrupting the membrane capacitance, and releasing interstitial fluid into the extracellular space. ^16, 37^ Overall, our biological assessment provides preliminary insight into underlying biomolecular changes in spheroids that may contribute to observed changes in spheroid opacity in response to radiation. However, this study was conducted using a single HNC cell line, which may limit generalizability, as radiation sensitivity can differ across cancer types. Future studies incorporating additional HNC subtypes and other cancer models will be important for broader validation. Nonetheless, our findings serve as a proof-of-concept for the application of rapid and minimally invasive LOC-EIS in assessing therapy response in 3D tumor models.

## Conclusion

Advancing LOC technologies for biological assessment can significantly lower barriers to developing and evaluating novel cancer therapies. The integration of label-free and non-invasive methods, such as EIS, which leverages cellular dielectric properties, offers potential in this direction. Incorporating EIS within a LOC enabled a quantitative assessment of radiotherapy-induced changes in individual 3D spheroids. This approach holds potential for design extension that may further enhance sensitivity and enable real-time tracking of spheroid degradation through its dielectric properties. Overall, our findings support the broader application of LOC-based EIS toward detecting therapy response in physiologically relevant 3D models, highlighting its promise in cancer research and drug development.

## Acknowledgements

This work was funded by the National Science Foundation NSF CAREER ECCS (2046037) to Leyla Esfandiari. Fabrication and development of the LOC prototype were supported by Colleen Arrasmith. Graphics were created using BioRender.com.

## Author Contributions

G.M., M.S. (Sheth), and L.E. conceived and designed the study. G.M., M.S. (Sheth), M.S. (Sharma), S.K., and M.L. performed the experiments. Data was curated and formally analyzed by G.M., M.S. (Sharma), and S.K. The experimental findings were validated and interpreted by G.M., M.S. (Sheth), M.S. (Sharma), S.K., and L.E. Figures and the original draft were prepared by G.M. The manuscript was edited by M.S. (Sheth), SK, T.W.D., V.T., and L.E. Supervision, funding acquisition, and project administration were provided by L.E. All authors reviewed and approved the final version of the manuscript.

## Ethics Approval

Ethics approval is not required.

