## Supporting Information for "Label-free monitoring of therapy response in 3D spheroids using lab-on-a-chip impedance spectroscopy"

| | 10kHz | 20kHz | 40kHz | 60kHz | 80kHz | 100kHz |
|---|---|---|---|---|---|---|
| Experiment 1 | | * | * | * | * | * |
| Experiment 2 | | | | * | ** | ** |
| Experiment 3 | | ** | *** | **** | **** | **** |

**Supplementary table 1.** Distinction of spheroid size by low-frequency impedance measurement replicate experiments. Table depicts at which evaluated discrete frequencies significant differences were found between the spheroids seeded with 2000 and 4000 cells.

| | 60kHz | 80kHz | 100kHz | 200kHz | 300kHz | 400kHz | 500kHz | 600kHz | 700kHz | 800kHz | 900kHz | 1MHz | 2MHz | 3MHz | 4MHz | 6MHz | 8MHz |
|---|---|---|---|---|---|---|---|---|---|---|---|---|---|---|---|---|---|
| Experiment 1 | | | | | | * | ** | ** | ** | ** | ** | | * | ** | * | | |
| Experiment 2 | | | | | * | * | ** | ** | ** | ** | ** | * | ** | ** | | | |
| Experiment 3 | | | | * | ** | ** | ** | ** | * | * | ** | ** | * | * | | | |

**Supplementary table 2.** Replicate experiments for impedance measurements of 3D spheroids 3 days after radiation treatment. Table depicts at which evaluated discrete frequencies significant differences were found between the control and spheroids irradiated at 4Gy.

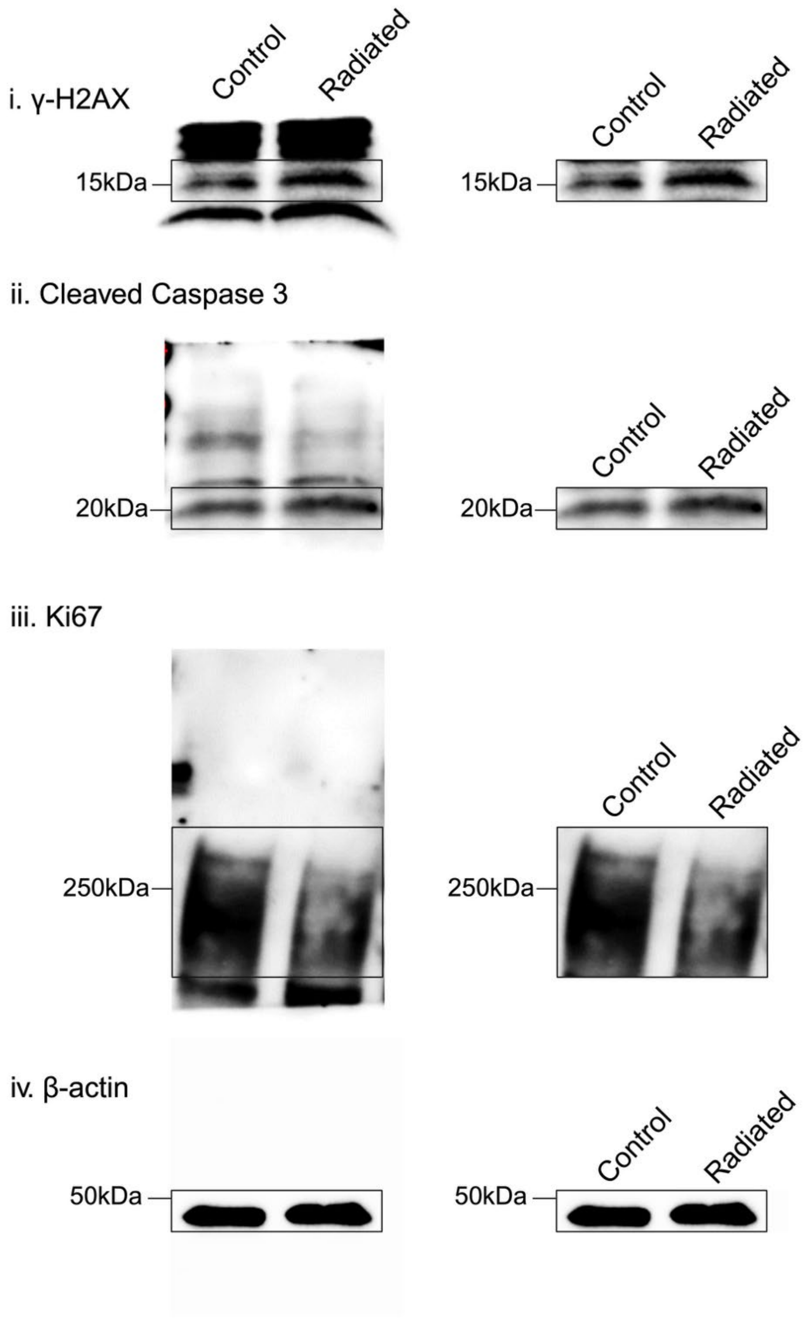


**Supplementary figure 1.** Raw western blot data for the evaluated biomarkers related to radiation response that may influence changing spheroid opacity.
